# Type 2 diabetes remodels collateral circulation and promotes leukocyte adhesion following ischemic stroke

**DOI:** 10.1101/2024.10.23.619748

**Authors:** Yoshimichi Sato, Yuandong Li, Yuya Kato, Atsushi Kanoke, Jennifer Y Sun, Yasuo Nishijima, Ruikang K. Wang, Michael Stryker, Hidenori Endo, Jialing Liu

## Abstract

Type 2 diabetes mellitus (T2DM) is associated with impaired leptomeningeal collateral compensation and poor stroke outcome. Neutrophils tethering and rolling on endothelium after stroke can also independently reduce flow velocity. However, the chronology and topological changes in collateral circulation in T2DM is not yet defined. Here, we describe the spatial and temporal blood flow dynamics and vessel remodeling in pial arteries and veins and leukocyte-endothelial adhesion following middle cerebral artery (MCA) stroke using two-photon microscopy in awake control and T2DM mice. Relative to control mice prior to stroke, T2DM mice already exhibited smaller pial vessels with reduced flow velocity. Following stroke, T2DM mice displayed persistently reduced blood flow in pial arteries and veins, resulting in a poor recovery of downstream penetrating arterial flow and a sustained deficit in microvascular flow. There was also persistent increase of leukocyte adhesion to the endothelium of veins, coincided with elevated neutrophils infiltration into brain parenchyma in T2DM mice compared to control mice after stroke. Our data suggest that T2DM-induced increase in chronic inflammation may contribute to the remodeling of leptomeningeal collateral circulation and the observed hemodynamics deficiency that potentiates poor stroke outcome.

**Highlights:**

- Blood flow and leukocyte imaging in awake mice by two-photon microscopy before and after stroke under physiological conditions
- T2DM induces collateral remodeling prior to stroke
- T2DM reduces blood flow and impedes recovery in pial arteries and veins after ischemic stroke
- Poor recovery of penetrating arterial flow and sustained deficit in microvascular flow after ischemic stroke in T2DM mice
- T2DM increases persistent leukocyte adhesion to endothelium of veins and elevates neutrophils infiltration into the brain parenchyma after ischemic stroke.

## 1. Introduction

Stroke is a highly prevalent and debilitating neurological disorder that is associated with significant rates of morbidity, disability, mortality, and recurrence. Cerebral collateralization directly contributes to the cerebrovascular reserve capacity and may potentially improve the hemodynamics after cerebrovascular occlusive disease like an ischemic stroke ^1, 2^. Established clinical studies have long reported that the status of the cerebral collateral circulation alters the risk and outcome of ischemic stroke. Type 2 diabetes mellitus (T2DM) is a major risk factor of ischemic stroke and is associated with an increased risk of long-term functional deficits after stroke ^3–5^. Patients with T2DM are associated with poor collateral status during acute ischemic stroke, which can drive rapid progression of the ischemic core ^6–8^. Impaired development of coronary collateral vessels was reported in patients with metabolic syndrome, indicating the chronic effect of metabolic disease on collateral remodeling ^9^. Consistent with the poor perfusion status, patients with acute ischemic stroke who present with acute hyperglycemia or diabetes are more likely to have a less favorable clinical outcome and 90-day mortality, regardless of recanalization method^10^. While established experimental evidence suggests that T2DM is associated with impaired leptomeningeal collateral compensation during MCA stroke and during mild ischemia ^11, 12^, the mechanisms underlying the poor collateral status in stroke patients with T2DM remains incompletely understood.

Neutrophils not only are implicated in the major processes that cause ischemic stroke, including thrombosis and atherosclerosis, they also contribute to ischemic brain injury ^13^. Activation of sympathetic nervous system during ischemic stroke leads to the release of catecholamine, which in turn stimulates spleen to release additional neutrophils into periphery that later migrated to the brain ^14^. Among the first responders after ischemic stroke, neutrophils enter into the ischemic brain regions through cerebral vessels, choroid plexus and subarachnoid spaces within hours ^15^ and promote neurovascular unit disruption, cerebral edema, and brain injury ^16^. Early increase of neutrophils plays a prominent role in instigating a larger stroke size and worse clinical outcomes among stroke patients with T2DM compared to patients without T2DM, increasing morbidity and mortality ^17^ ^18^. Neutrophils can also promote thrombosis via NETosis and interact with platelets post stroke. Lastly, neutrophils engaging in tethering and rolling on endothelium after stroke can directly reduce flow velocity.

The effect of T2DM on neutrophil adhesion, hemodynamics and their progression during ischemic stroke remains elusive. To test the hypothesis that T2DM-worsened inflammation would affect leukocyte adhesion on endothelium and impact blood perfusion after stroke, we examined the blood flow dynamics in pial arteries and veins and the spatial and temporal changes of the leukocyte adhesion following MCA stroke in normal and T2DM mice using two-photon microscopy in awake mice. The number of neutrophils infiltrated into parenchyma during acute stroke was quantified by flow cytometry. We additionally determined the effect of T2DM in downstream penetrating arterioles and capillary vessels by optical coherence tomography angiography.

## 2. Materials and methods

### 2.1. Animals and housing

All animal experiments were conducted in accordance with the *Guide for Care and Use of Laboratory Animals, and reported in compliance with the Animals in Research: Reporting In Vivo Experiments (ARRIVE) guidelines* ^19, 20^, and were approved by the San Francisco Veterans Affairs Medical Center Institutional Animal Care and Use Committee. Male db/+ and db/db mice (16–20 weeks old, The Jackson Laboratory) were housed 4 per cage on a 12 h dark/light cycle with access to food and water prior to experimental procedures.

### 2.2. Animal randomization for treatment and blinding of identity

Mice were coded at the beginning of the experiment and randomized into treatment groups. Although the mouse strain was apparent to investigators performing in vivo procedures owing to difference in body weight, the identity of each mouse subject was blinded to investigators who conducted the experiments and data analysis. Data from 57 db/db and 52 db/+ mice were reported in this study. Sample sizes for individual experiments are listed in the figure legends. Approximately 20% db/+ and 50% db/db mice with cranial windows of insufficient quality for 2p and OCT imaging were excluded before the induction of experimental stroke. The low successful rate of the latter mice was attributed to the increased bleeding tendency and poor wound healing capacity. Additional 5 db/+ and 9 db/db mice did not contribute to the final data set in two-photon and OCT imaging owing to ischemic injury-related mortality compounded by repeated imaging before reaching the final time point. Mortality rate due to stroke in mice used in flow cytometry experiment was less than 10%. Lastly, data from 2 db/+ and 8 db/db mice and from 1 db/+ and 7 db/db mice were excluded from the penetrating arterial flow and microvascular velocimetry results due to insufficient MCA vessels imaged/quantified.

### 2.3. Cranial window implantation for imaging

Two weeks prior to two-photon imaging, mice were anesthetized and placed on a stereotaxic frame. An incision of the scalp in the midline was made to expose the skull, and a head-fixation metal plate with a hole in the center was fixed to the left side of the skull. A 6-mm-diameter craniotomy was performed with a dental drill on the parietal cortex 5 mm lateral and 1 mm posterior from the bregma. A circular piece of skull was carefully detached and replaced with a round coverslip glass 4 mm in diameter. The coverslip was sealed to the skull with 3 M VetBond (St. Paul, MN), cyanoacrylate glue and dental cement, producing a cranial window for high-resolution blood flow imaging. The pulled-back skin was sealed to the skull with 3 M VetBond. Rectal temperature was maintained within the range of 37 ± 0.5°C by the use of a heating blanket and rectal thermistor servo-loop throughout the procedure. Cranial window implantation for OCT imaging was similarly performed without fixation of the metal plate.

### 2.4 Experimental stroke

Ischemic stroke was induced by permanent left middle cerebral artery occlusion (MCAO) plus 60 min of left CCAO, then removal of the CCA occlusion using the distal MCAO method as previously described with modifications^11, 21^. Specifically, after ligation of the ipsilateral CCA, the distal M1 portion of left MCA, peripheral to the perforating arteries of the basal ganglia, was permanently occluded by electrocauterization and dissected. During the operation, blood pressure was monitored by an indirect blood pressure meter and rectal temperature was regulated within the range of 37 ± 0.5°C with a heating blanket and rectal thermistor servo-loop throughout the procedure. Sham-operated mice did not receive occlusion of either the MCA or CCA.

### 2.5. Longitudinal two-photon imaging and blood flow analysis

To label plasma and leukocytes/platelets, respectively, Texas Red dextran (40 μl, 2.5%, MW=70,000 kDA, Thermo Fisher Scientific) and Rhodamine 6G (100 μl, 1mg/ml in 0.9% saline, Acros Organics, Pure) were injected into the jugular vein under isoflurane anesthesia before imaging ^22^. Animal was fixed into a customized rotary stabilization bar for high-resolution in vivo imaging using a resonant-galvo scanning, two-photon microscope (Neurolabware, Los Angeles, CA, USA), during which the acquisition was controlled by MATLAB-based Scanbox (Neurolabware) software. The light source is a mode-locked Ti: sapphire laser (Coherent Chameleon Ultra II) with excitation wavelength of 840 nm. Excitation power measured at the back aperture of the objective was typically less than 50 mW and adjusted in each session to achieve similar levels of fluorescence. Emission light was filtered by emission filters (525/70 and 610/75 nm) and measured by two independent photomultiplier tubes (PMTs) (Hamamatsu R3896), referred to as red and green channels. Images were acquired with a Nikon 16X water immersion objective (NA =0.8, 3 mm working distance) (Figure 1A). Using the surface vasculature and epifluorescence imaging, the same imaging areas were identified between session. For each imaging session, 20-30 ROIs was imaged. Baseline imaging sessions began 2-3 weeks after cranial window surgeries, to ensure surgery-induced inflammation and gliosis has subsided post cranial window implantation. Repeated imaging was performed at four time points: baseline prior to stroke (labeled as Base), one hour after the reperfusion of ipsilateral CCA (as 1h), one day or seven days after MCAO (as 1d and 7d). At each imaging location, blood vessels were imaged at a depth of 100 - 300 micron below the pial surface. Full-field images at the frame rate of 15.5Hz (512 lines with a resonant mirror at 8 kHz) were collected, and then 2s line-scanning along the central axis of a selected blood vessel was performed at 8kHz to calculate blood flow velocity. Arterial branches imaged were classified according to the proximity to occlusion site, with most distal MCA branches assigned as segment 1 (S1) and the most proximal ones as segment 3 (S3) as shown in the cranial window view (Figure 1B). Vein branches imaged were also classified according to the proximity to the sinus they drain out. The most distal veins draining out to the Superior Sagittal Sinus (SSS) were labeled as anterior cerebral artery (ACA) vein branches, in which the distal ones were assigned as ACA1 (A1) and the proximal ones as ACA 2 (A2). Similarly, the veins draining out to the Sylvian vein or superficial middle cerebral vein were labeled as MCA vein branches, in which the distal ones were assigned as MCA1 (M1) and the proximal ones as MCA 2 (M2). Image stacks of 200 frames (lasting for 12.5 seconds with the full-field 512 line images) were captured to calculate leukocyte-platelets adhesion.

**Figure 1.**
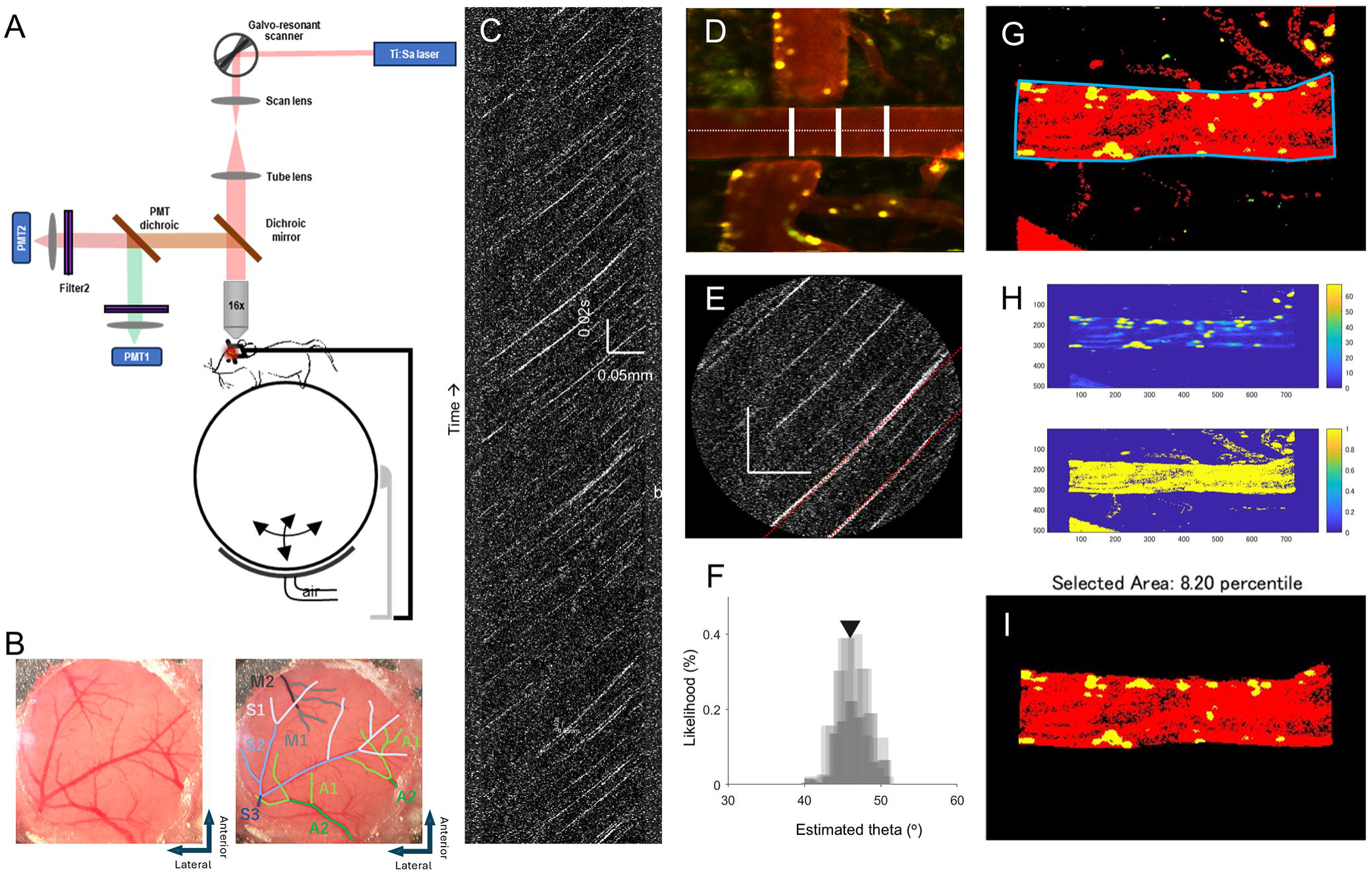
The schematic diagram of the two-photon system and methods to determine vascular velocity, diameter and the area of adherent leukocytes. (A) The schematic of the custom-designed two-photon system. Mouse was fixed to a customized rotational stage with a head plate in an awake state, with limbs freely moving on an air-floated Styrofoam ball. (B) An original image of the cranial window accompanied by a replica with showing labeled pial arteries in 3 segments (S1: distal, S2: middle, S3: proximal) and pial veins (draining to superior sagittal sinus: A1: distal ACA vein branch, A2: proximal ACA vein branch; draining to Sylvian fissure: M1: distal MCA vein branch, M2: proximal MCA vein branch). (C) A two-photon kymograph of the line-scan. Blood flow velocity was measured using line-scan images along the central axis of the cortical arteriole. (D) Measurement of vessel diameter. Blood vessel diameter was determined by measurements taken at three different cross-sections of the vessel. (E) Raw line scan images flattened by temporal demeaning and Sobel filtering for velocity calculation. (F) Radon Transform of the linear RBCs streak. (G) To calculate the area of adherent leukocyte on vessel wall over the cross section of the blood vessel, ROI were manually curated to identify the cross section area of the veins of interest. (H) Adaptive thresholding was applied to each frame of the green channel to identify the leukocyte adhesion (upper row) and global thresholding was applied to the red channel to identify the overall blood vessel area (lower row). (I) The leukocyte area/cross section vessel area ratio was calculated within the identified region of interest.

### 2.6. Data Analysis of Two-photon imaging

Two-photon images were analyzed in MATLAB using customized code blinded to experimental condition and imaging time point. Blood flow velocity was measured using line-scan images along the central axis (Figure 1C). The analysis area was defined by user using the full-filed image stacks, and blood vessel diameter was determined by measurements taken at 3 different cross-sections of the vessel (with the variation less than 1%) (Figure 1D). The first 0.1 second (5% frames) was discarded due to potential motion of the scan head. Raw images are flattened and denoised by temporal demeaning and Sobel filtering, and the angle of the linear RBCs streaks were determined using iterative Radon transform (Figure 1E, F) ^23, 24^. Velocity is then calculated by scaling the tangent of this angle with the temporal (Δ time) and spatial (Δdistance) resolution of the images, using the following equation: Velocity= tan(θ) * Δ time/Δ distance. Finally, RBC flux was calculated from measurements of blood flow velocity and vessel lumen diameter based on previous work by Kleinfeld’s group ^25^ using the following equation: Flux=π/8 × velocity × diameter^2^.

The relative amount of leukocyte adhesion to vessel wall varies by the vessel diameter and expressed as the percent area of adherent leukocyte on vessel wall over the cross section of the blood vessel, calculated from image stacks of 200 frames converted from the recorded video. ROI was manually curated according to the veins of interest in order to exclude signals from any other vessel (Figure 1G). The first 0.1 second (less than 1% frames) was discarded due to potential motion of the scan head. Motion correction was then applied using a recursive imaging alignment based on a MATLAB code modified from Scanbox. Global thresholding was applied to the red channel to identify the overall blood vessel area, and adaptive thresholding was applied to each frame of the green channel to identify the leukocyte adhesion on vessel wall (Figure 1H). The leukocyte adhesion ratio was then calculated by the identified stalled leukocyte adhesion area normalized by the blood vessel area (Figure 1I). On average 15 artery and 8 vein locations were imaged at each time point for each mouse.

### 2.7 Blood flow imaging via optical coherence tomography angiography (OCTA)

Cortical penetrating arterial microvascular flow dynamics were quantified by a custom OCT system as previously described ^11, 12, 21^ (Supplemental Figure 1A).

### 2.8. Brain Mononuclear Cell isolation

At 1, 3 or 7 days after MCAO or 1 day after sham surgery, 3 ipsilateral forebrain hemispheres were harvested and pooled for each time point per genotype after cardiac perfusion with heparinized GKN buffer (8 g/L NaCl, 0.4 g/L KCl, 1.41 g/L Na2HPO4, 0.6 g/L NaH2PO4, 2 g/L D(+) glucose, pH 7.4) as previously described ^30^. The tissues were digested in NOSE buffer (4.0 g/L MgCl2, 2.55 g/L CaCl2, 3.73 g/L KCl, 8.95 g/L NaCl, pH 6-7) supplemented with 200 U/ml DNase I (Catalog No. D5025-150KU, Sigma-Aldrich, St. Louis, MO) and 0.2 mg/ml collagenase type I (Catalog No. L5004196, Worthington Biochemical, Lakewood, NJ) at 37°C for 30 min. Thoroughly washed cells were subjected to an isotonic Percoll gradient (90% Percoll, Catalog No. 17-0891-01, GE Healthcare Life Sciences, Chicago, IL) by suspending cells in 4 mL of 1.03g/mL Percoll in GKN buffer/0.2%BSA and overlaying cell suspension on 4 mL of 1.095 g/mL Percoll in PBS/0.2%BSA. After centrifugation at 900xg for 20 min at RT, brain mononuclear cells were collected from the interface, washed, and processed further for FACS staining and analysis.

### 2.9. Fluorescence-activated cell sorting (FACS)

Brain mononuclear cells were suspended in 100 µL FACS buffer and blocked with 1µL anti-mouse CD16/CD32 (clone 93, Catalog No. 14-0161-85, eBioscience) for 20 min. The cells were immunostained for 30 min on ice with 1µL of each of the following antibodies against: CD11b (clone M1/70, PerCP 5.5, Catalog No. 45-0112-82, eBioscience), CD45 (clone 30-F11, FITC, Catalog No. 11-0451-81, eBioscience), Ly6G Gr-1 (clone 1A8-Ly6G, PE-eFluor 610, Catalog No. AB_2574679, eBioscience), and fixable viability dye (eFluor 506, Catalog No. 65-0866-14, eBioscience). 50 µL of Absolute counting beads (Catalog No. C36950, Life Technologies, Carlsbad, CA) were used for cell counting. After washing, the cells were resuspended in 100 µL of FACS buffer and fixed with 100 µL 2% PFA. Data were acquired using a BD FACS ARIAIII and analyzed using FlowJo v10 software (FlowJo).

### 2.10. Statistical analysis for two-photon microscopy and OCT

Data were expressed as Mean ± SD. The Kolmogorov Smirnov normality test was used to determine whether data were normally distributed. Data were analyzed by two-way repeated measure ANOVA for temporal and genotype differences or by two-sided t-test followed by Bonferroni corrections for multiple comparisons using Prism 10 (GraphPad Software Inc., San Diego, CA). FACS data were analyzed at each time point using the Mann-Whitney U test individually. Priori sample size calculations were performed based on our prior studies with this stroke model for two-photon microscopy and for OCT angiography. P values less than 0.05 were considered significant.

## 3. Results

### 3.1. Pial veins and distal pial arteries of diabetic mice are smaller in size and slower in flow velocity prior to stroke

To determine how native collaterals affect retrograde and anterograde blood flow spatially and temporally in db/+ and db/db mice, wide field two-photon imaging was employed to quantify flow over distal MCAs (Figure 2A). At baseline, a decreasing gradient in flow velocity was detected from S3 to S1 in both genotypes (Figure 2B). A decreasing gradient was also detected in the diameters from S3 to S1 in both genotypes. Flow velocity is much slower in pial veins compared to pial arteries, despite the comparable size between pial veins and S3 segments of the pial arteries. In addition, the velocity, diameter and flux of pial veins of db/db mice was significantly lower compared to those of db/+ mice. In contrast, db/+ mice had a significantly higher velocity and greater diameter in distal (S1 and S2) MCA branches and veins. It suggests that T2DM chronically remodels the leptomeningeal vessels prior to stroke.

**Figure 2.**
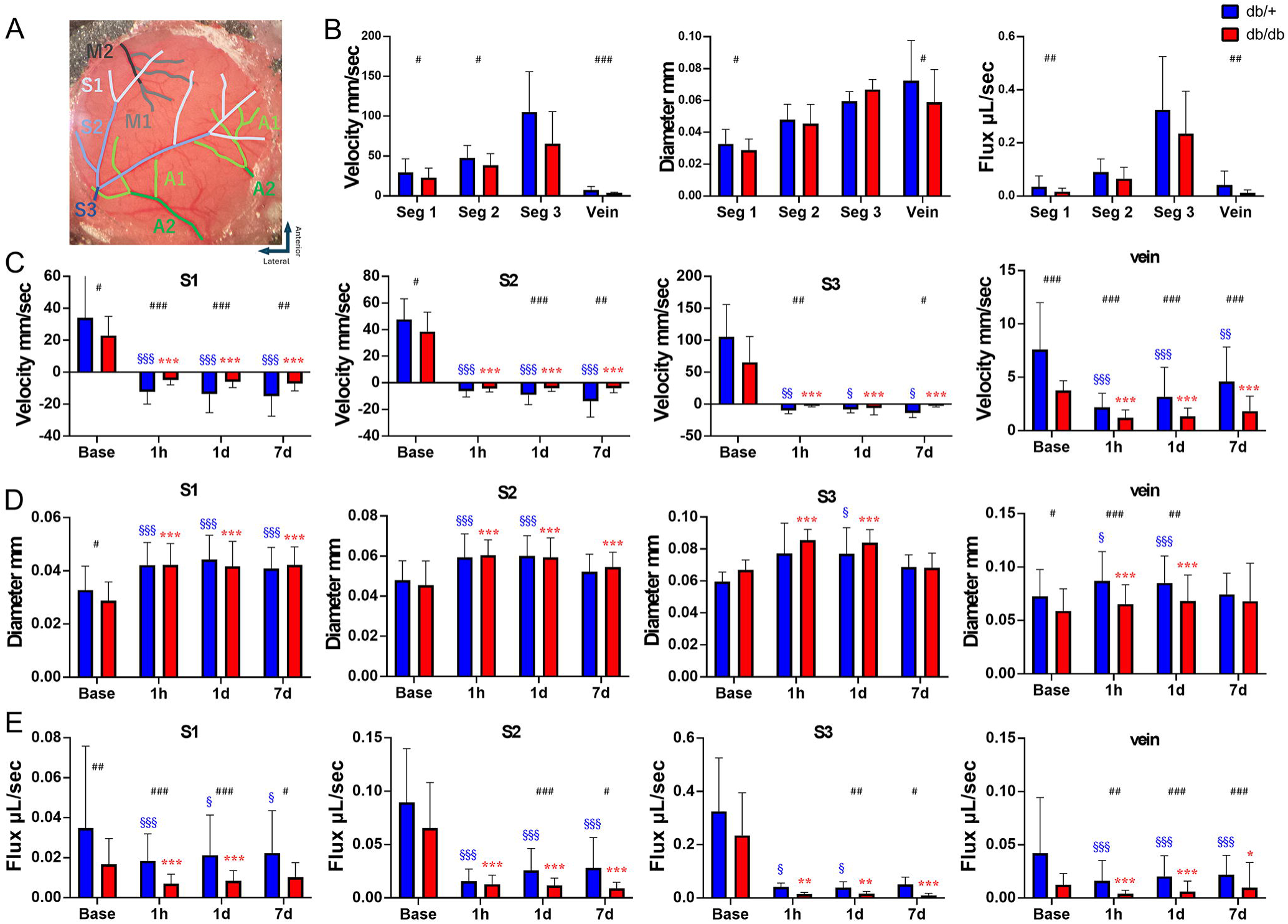
Diabetic mice have a poor collateral flow in the ischemic territory after stroke. (A) Labeled vascular segment map within the cranial window. (B) Base line value of velocity, diameter and flux in artery and veins. db/db mice have significantly slower velocity and smaller diameter in S1 and veins. As a result, the flux in S1 and vein in db/db mice are significantly smaller compared to db/+ mice. (C) Flow reconstruction was observed immediately after MCAO. Flow direction changes were observed in S1-S3 after MCAO, with the retrograde flow from the ACA towards the MCA. db/db mice had significantly slower velocity in all segments and at most timepoints. The flow velocity of MCA branches and veins progressively improved in db/+ mice toward 7d after MCAO. In contrast, no improvement was seen in db/db mice in any vessels within the cranial window. (D) Dilation of pial arteries after MCAO was most robust and persistent in distal MCAs where the retrograde collateral flow began for both type of mice. In the most proximal segment (S3), the dilation declined at 7d in both genotypes. A significantly bigger diameter at baseline or a greater extent of dilation was seen in the pial veins of the db/+ mice compared to those of db/db ones till 1d after MCAO. (E) The flux dramatically decreased after MCAO. db/db mice had a smaller flux compared to db/+ mice in all segments and at most of the timepoints. The flux of MCA branches progressively improved in db/+ mice toward 7d after MCAO but not in db/db mice. Time point vs. baseline: §, §§, §§§P < 0.05, 0.01, 0.005 (db/+); *, **, ***P < 0.05, 0.01, 0.005 (db/db). Strain difference: #, ##, ###P < 0.05, 0.01, 0.005. N = 11 db/+, 9 db/db.

### 3.2. Diabetic mice have a poor collateral flow in the ischemic territory after stroke

A robust reversal flow of MCA branches was observed after MCAO and lasted till 7d, the longest time point in our study (Figure 2C). Decreasing gradient in flow velocity was detected from S1 to S3 in both genotypes. The flow velocity of the MCA branches slowed down more dramatically in db/db mice in veins and in almost all segments of arteries and for almost all time points after stroke. The flow velocity of MCA branches showed varying extent of progressive recovery in db/+ mice toward 7d after MCAO, in stark contrast to almost no recovery in the db/db mice. Pial arterioles and veins of both genotype dilated after stroke at most locations during hyper acute stroke phase (Figure 2C). Dilation of S3 was terminated at 7d after MCAO, possibly due to infarct expansion. Similar to baseline, db/+ mice had a significantly larger diameter of pial veins after MCAO and lasted till 1d. The flux dramatically decreased after MCAO and progressively recovered in db/+ mice toward 7d but not in db/db mice. db/+ mice also had a significantly higher flux in most of the arterial segments and veins during most of the time points (Figure 2D). These results suggest that the db/+ mice had a better collateral flow of the ischemic territory compared to db/db mice, which also recovered more favorably after MCAO.

### 3.3. MCA branches with anterograde flow have a miserable perfusion and do not recover after stroke

Some distal MCA branches (S1, S2) maintained an anterograde flow after permanent MCAO (Figure 3A-C). These branches usually exhibited slower velocity compared to the parent arteries they branched out (Figure 3A). Among these MCA branches, a dramatic decrease in velocity and flux in both genotypes was observed and they did not recover till at least 7d after MCAO (Figure 3D). The velocity and flux of S1 and S2 with the anterograde flow were significantly lower compared to those MCA branches with retrograde flow in the same segment territory in both genotypes. These results suggest that MCA branches without direct collateral from the ACA branches have a miserable collateral flow and a low ability to recover after stroke.

**Figure 3.**
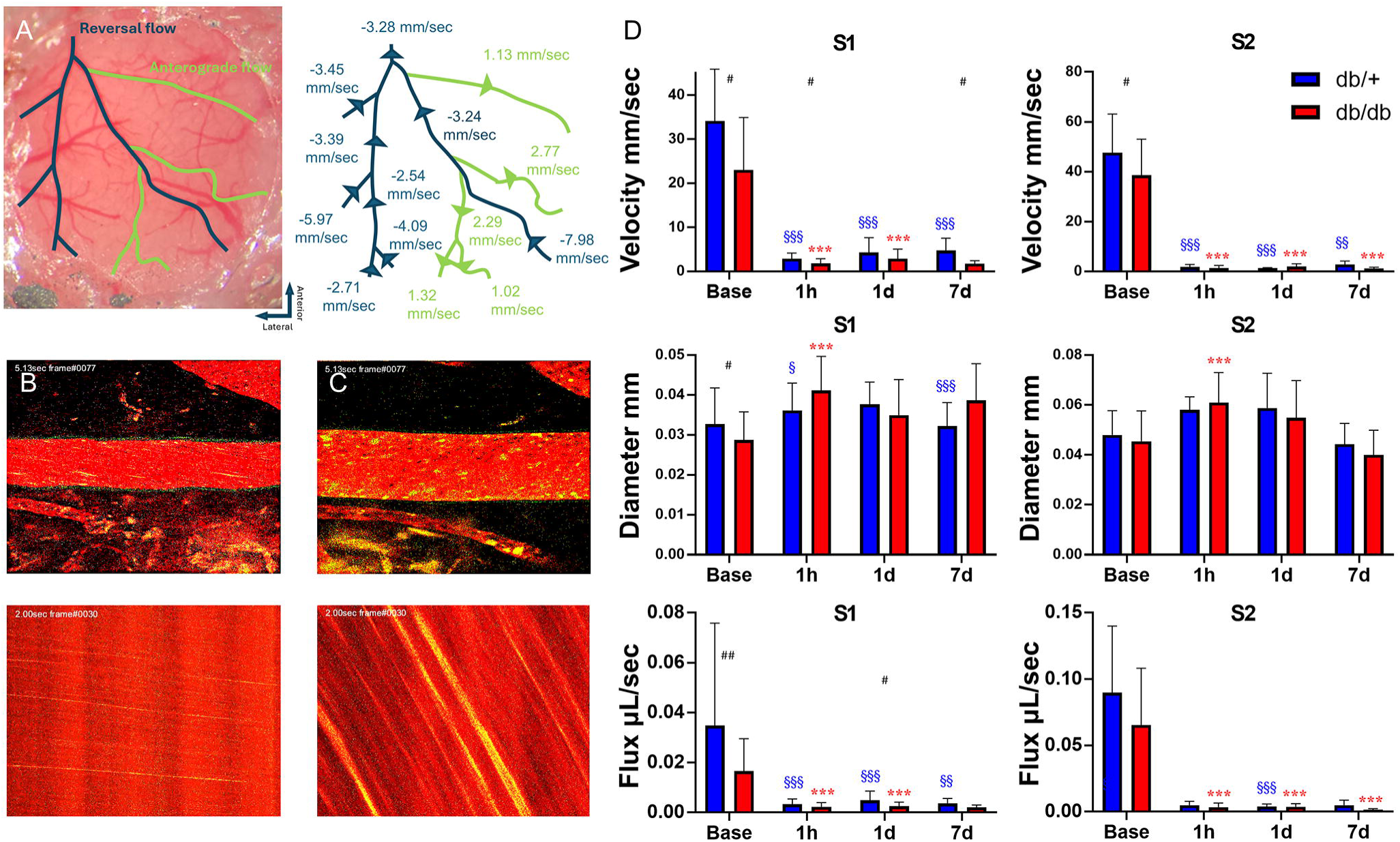
MCA branches with anterograde flow have a miserable perfusion and do not recover after stroke. (A) Labeled vascular map with flow directions and the value of flux at various locations within the cranial window, in which the retrograde flow is labeled in blue and anterograde flow in green. The flux of S1 and S2 with anterograde flow is reduced compared to their parent branches with retrograde flow. (B, C) Examples of two-photon microscopy images of S1 in db/+ mice at base line (B) and after MCAO showing anterograde flow (C). Dilation of the pial artery and a decrease of velocity were observed. (D) Velocity, diameter and flux of S1 and S2 with anterograde flow after MCAO. Significant decrease of velocity and flux was observed at all time points with little or no improvement in both strains. Slight dilation was also observed but gradually returned to base line till 7d after MCAO. Time point vs. baseline: §, §§, §§§ P < 0.05, 0.01, 0.005 (db/+); *, **, *** P < 0.05, 0.01, 0.005 (db/db). Strain difference: #, ##, ### P < 0.05, 0.01, 0.005. N = 11 db/+, 9 db/db.

### 3.4. Diabetic mice show greater leukocyte adhesion in the veins after MCAO

Diabetic db/db mice showed an increase of leukocyte adhesion in the veins in 1d and lasted till at least 7d after MCAO, in contrast to the gradual reduction seen in the veins of control db/+ mic (Figure 4A, B). To test the hypothesis that leukocyte adhesion tends to occur after stroke in vessels with smaller diameter and slower velocity, we compared leukocyte adhesion in the distal (A1) and proximal (A2) segment of the veins draining into SSS. Two-way ANOVA found differences in the main effect of vein segment in velocity (p<0.01), diameter (p<0.001), and flux (p<0.0001), showing that the A1 segment is smaller and has slower velocity. However, the difference in leukocyte adhesion between A1 and A2 segments did not reach statistical significance (p = 0.13). There was also a main effect of T2DM on velocity (p<0.0001), diameter (p<0.0001), flux (p<0.0001), and leukocyte adhesion (p<0.01). Specifically, db/db mice had a slower velocity, diameter and flux in A2 compared to db/+ mice prior to stroke (Figure 4C). The velocity of A1 and A2 was significant slower in both genotypes immediately after MCAO compared to baseline flow rate. The velocity of A2 segment in db/+ mice was not only significantly faster than that of the db/db mice but also progressively recovered toward 7d after MCAO, in contrast to the db/db mice. A1 veins of the db/+ mice dilated after MCAO compared to baseline in contrast to no dilation of the db/db veins, leading to greater flux and drainage of db/+ blood into the SSS. Significant increase of leukocyte adhesion was recognized mainly in A1 after MCAO compared to baseline and lasted till 7d in both genotypes. Genotype difference in leukocyte adhesion in either segment was lost due to smaller individual sample sizes compared to pooled sample size in Figure 4A.

**Figure 4.**
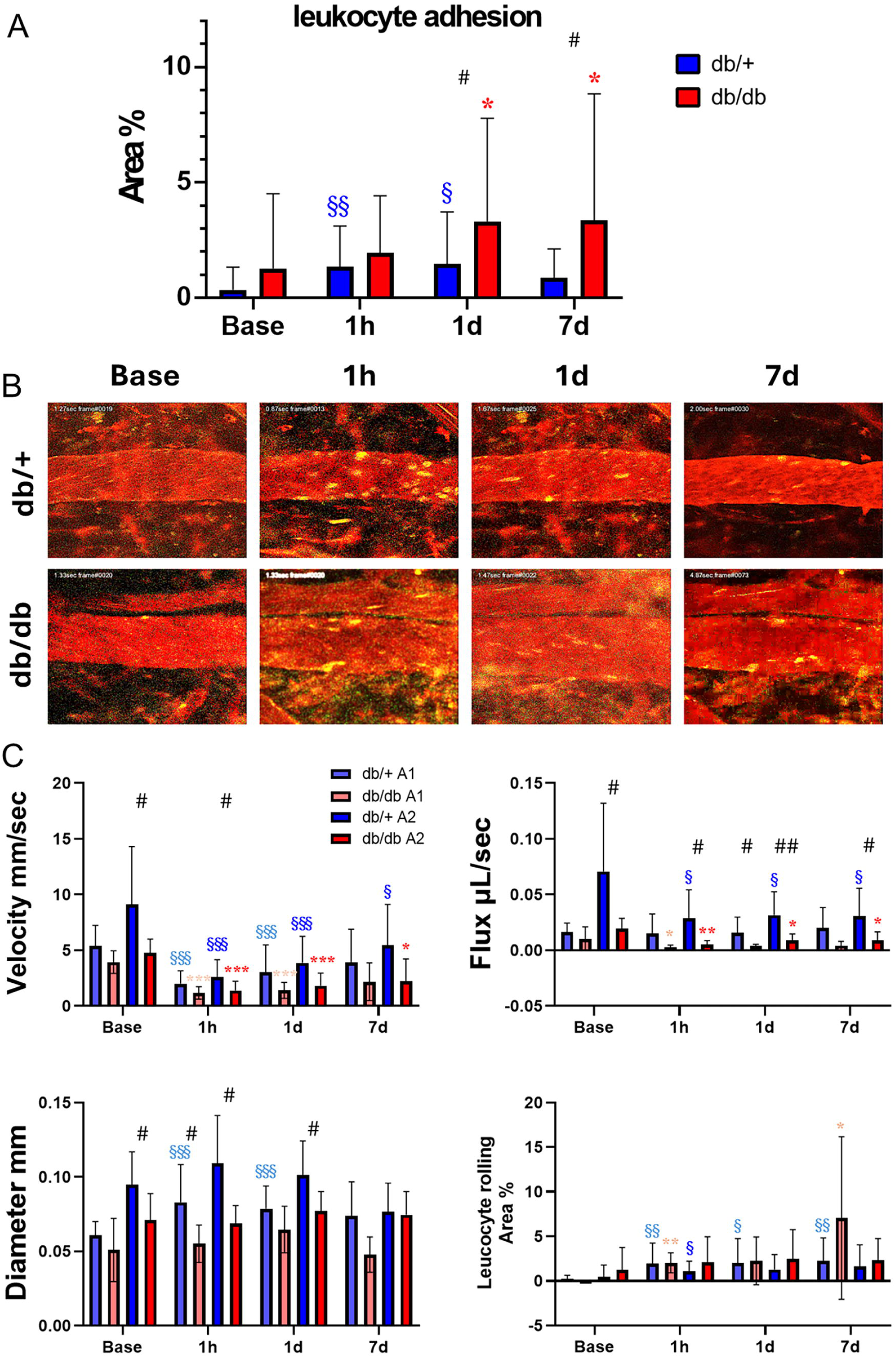
Diabetic mice show greater leukocyte adhesion in the veins after MCAO. (A) The leukocyte adhesion ratio in the veins of db/+ mice increased till 1d after MCAO in contrast to the persistent increase till at least 7d after MCAO in the db/db mice. Significant genotype difference was detected at 1d and 7d. (B) Two-photon image of the veins of db/+ and db/db mice. Leucocyte adhesion is observed immediately after MCAO and last till 7d in db/db mice. (C) Velocity, diameter, flux and leucocyte adhesion in each vein segments during and after MCAO. The flow velocity and flux of A2 segment proximal to the SSS has shown robust and progressive improvement at 7d after MCAO in db/+ mice, potentially contributing to the clearance of the leukocyte-platelet aggregates and reduction of adhesion. In contrast, db/db mice had larger increase of leukocyte adhesion in 7d but the genotype difference was not significant. While the analysis did not lead to significant statistical difference due to insufficient sample sizes, leukocyte adhesion tended to increase in smaller veins that are distal to the SSS with slower velocity. Time points vs. baseline: §, §§, §§§ P < 0.05, 0.01, 0.005 (db/+); *, **, *** P < 0.05, 0.01, 0.005 (db/db). Strain difference: #, ##, ### P < 0.05, 0.01, 0.005. N =11 db/+, 9 db/db.

### 3.5. Diabetic mice have more neutrophil infiltrated into brain parenchyma after stroke

To determine the consequence of increased leukocyte adhesion observed in the pial veins of diabetic mice after stroke, we quantified the number of cells infiltrating into brain parenchyma during acute stroke. We found that the diabetic mice had significantly increased neutrophils in the brain at baseline, as well as one and three days after stroke compared to control mice (5A, B). In both type of mice, the infiltrating neutrophils peaked at 1d after stroke and gradually declined over the course of one week (5C), in contrast to the delay increase of macrophages at 3-7 days after stroke (not shown).

### 3.6. Diabetic mice suffer from a greater reduction of penetrating anterior flow after stroke and poorer recovery

A robust reduction of the ACA and MCA PA flow was observed immediately after MCAO in both genotypes and the flow reduction was more dramatic in db/db mice (Figure 6A). The flow velocity of MCA PA branches recovered significantly in db/+ mice as early as one hour after CCA reperfusion, reaching at least 50% of the baseline velocity (Figure 6B). In contrast, db/db mice did not show a significant degree of recovery in both total MCA PA flow and PA velocity even after 1d. The reduction of ACA PA flow and velocity was mild immediately after MCAO and almost completely recovered at 1 h after CCA reperfusion. No significant genotype difference was observed in the ACA PA branches for total flow and velocity.

**Figure 5.**
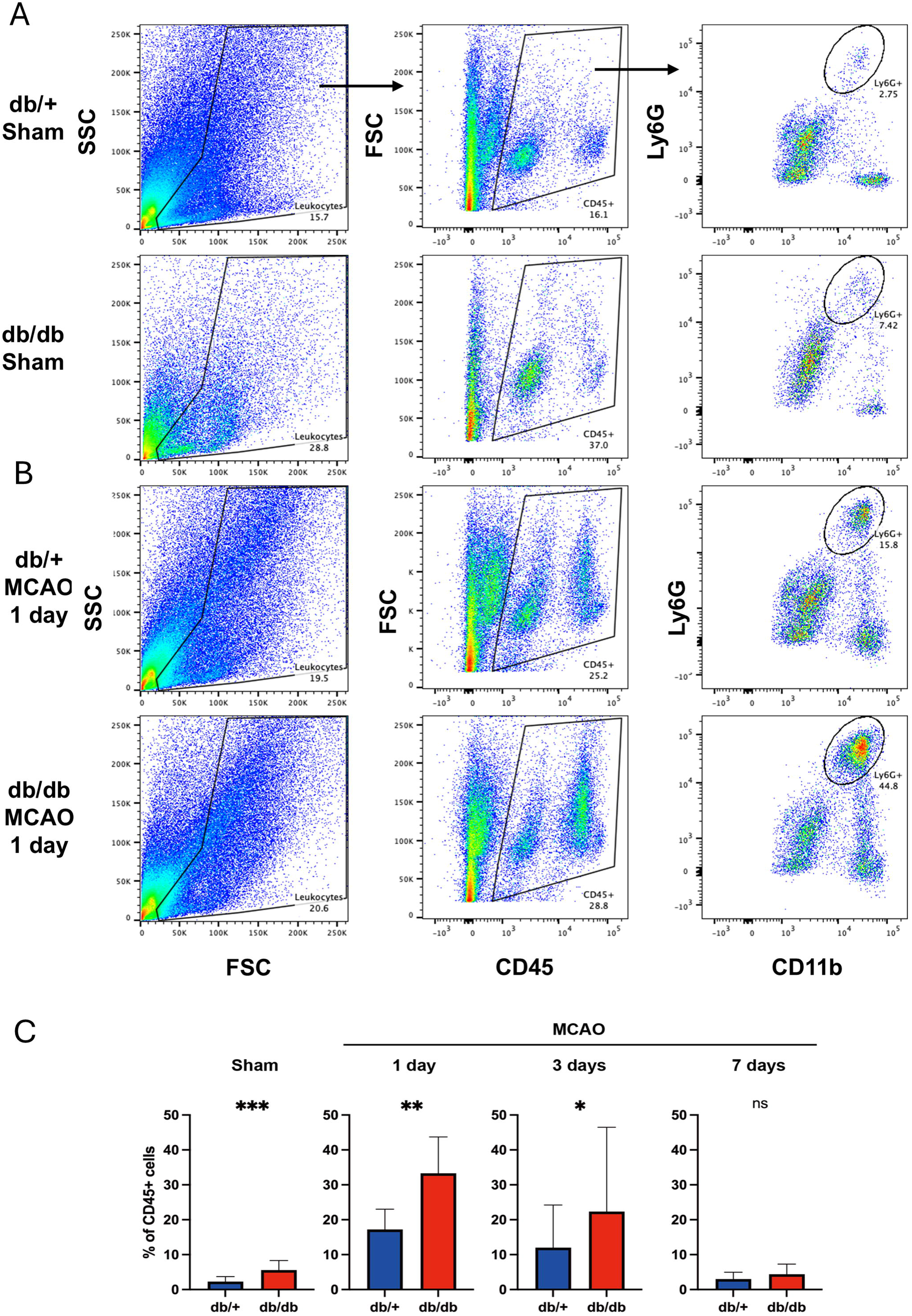
Diabetic mice have more neutrophil infiltrated into brain parenchyma after stroke. (A) Representative FACS gating scheme for brain leukocytes showing the proportion of infiltrated neutrophils (*CD45^+^/CD11b^hi^/Ly6G^hi^*) in the brains of the sham-operated groups of db/+ and db/db mice. (B) FACS plots showing the proportion of infiltrated neutrophils in the brain at 1 day after ischemic stroke in both db/+ and db/db mice. (C) Comparative results of neutrophil (Ly6G+CD11b+) proportions within CD45+ cells between the two groups at different time points: sham, 1 day, 3 days, and 7 days after ischemic stroke. The sample sizes are n=12 for Sham, n=10 for 1 day and 3 days, and n=8 for 7 days with equal number of mice between the two genotypes. ***p < 0.001, **p < 0.01, *p < 0.05. MCAO indicates Middle Cerebral Artery Occlusion.

**Figure 6.**
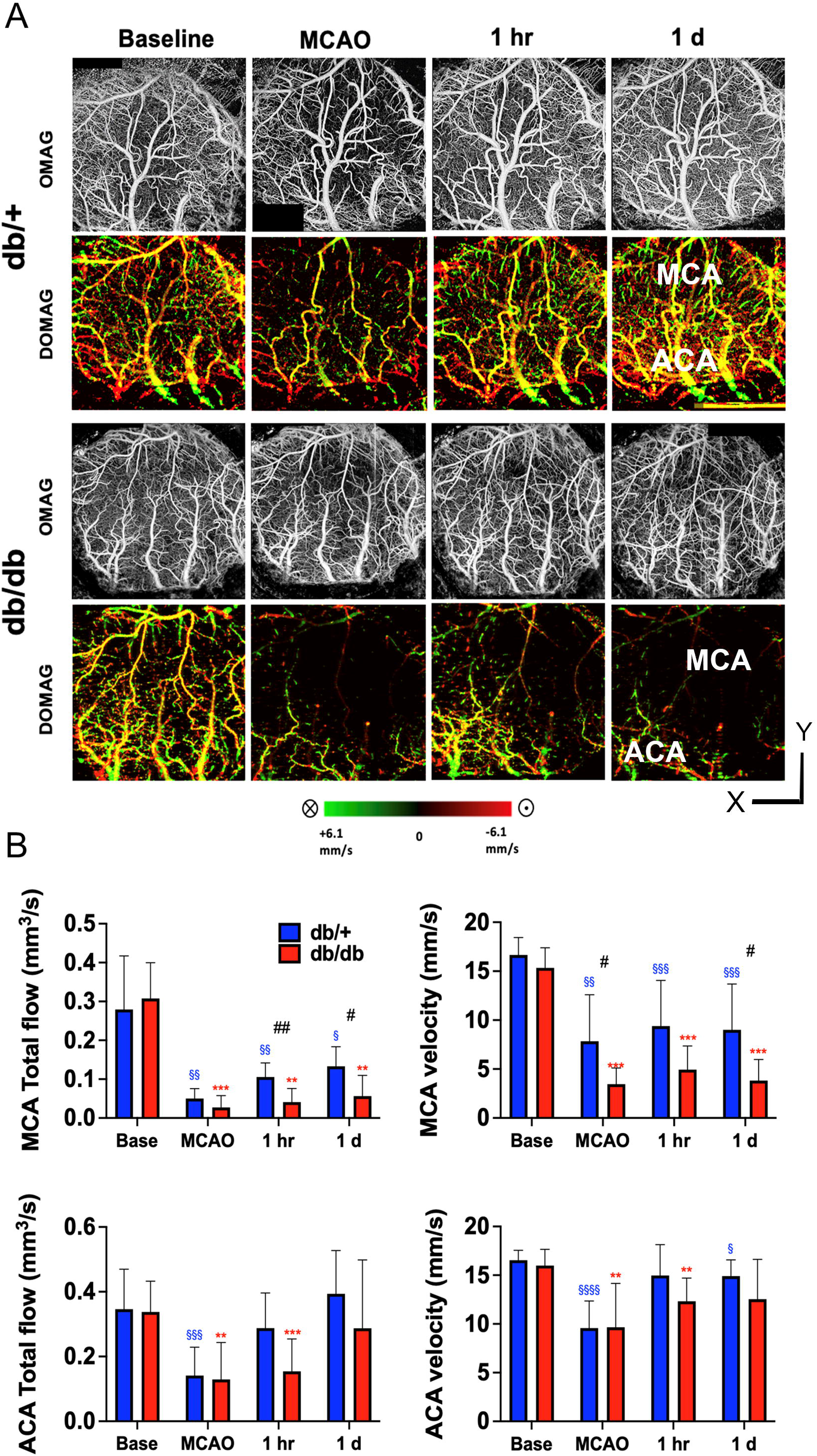
Diabetic mice suffer from a greater reduction of penetrating arterial flow and lesser recovery after stroke. (A) Representative *en face* maximal intensity projection of OMAG data (upper panel) and *en face* average intensity projection of 3D bidirectional DOMAG data (lower panel) within the 300-um-thick tissue slab below the cortical surface of db/+ (upper panels) and db/db (lower panels) mice showing dramatic reduction of penetrating arterial (PA) in both genotypes following MCAO. (B) PA flow was dramatically reduced immediately after MCAO in the MCA territory for both strains, although it was significantly recovered in db/+ mice at 1d compared to db/db mice. In contrast, the reduction of PA flow in the ACA territory was similar between genotypes and with much more robust recovery. Maximal MCA and ACA PA axial velocity was significantly higher in the db/+ mice compared to the db/db mice during the acute stage of MCAO. Time point vs. baseline: §, §§, §§§ P < 0.05, 0.01, 0.005 (db/+); *, **, *** P < 0.05, 0.01, 0.005 (db/db). Strain difference: #, ##, ### P < 0.05, 0.01, 0.005. N= 12 db/+, 6 db/db.

### 3.7. Diabetic mice display greater heterogeneity in capillary transit time after stroke compared to control mice

We next assessed and compared the mean and the spatial distribution of CTT temporally after stroke between db/+ mice and db/db mice via OCT imaging. The capillary MF maps obtained from OCTA velocimetry were shown after masking vessels with diameters larger than 15 µm (Figure 7A). The spatial distribution characteristics of the transit time, including mean CTT and CTTH were deduced from these MF signals for both the MCA and ACA region as described in the previous literature.^21^ Mean CTT and CTTH significantly increased after MCAO in db/db mice and persisted till 1d in the MCA and ACA territory (Figure 7B). db/+ mice showed no significant increase after MCAO. There was significant genotype difference in CTT and CTTH in the MCA territory and CTT in the ACA territory during MCAO. There was no significant strain difference in CTTH in the ACA territory. These findings suggest the db/db mice have a higher microvascular resistance and reduced oxygen extraction after MCAO compared to db/+ mice.

**Figure 7.**
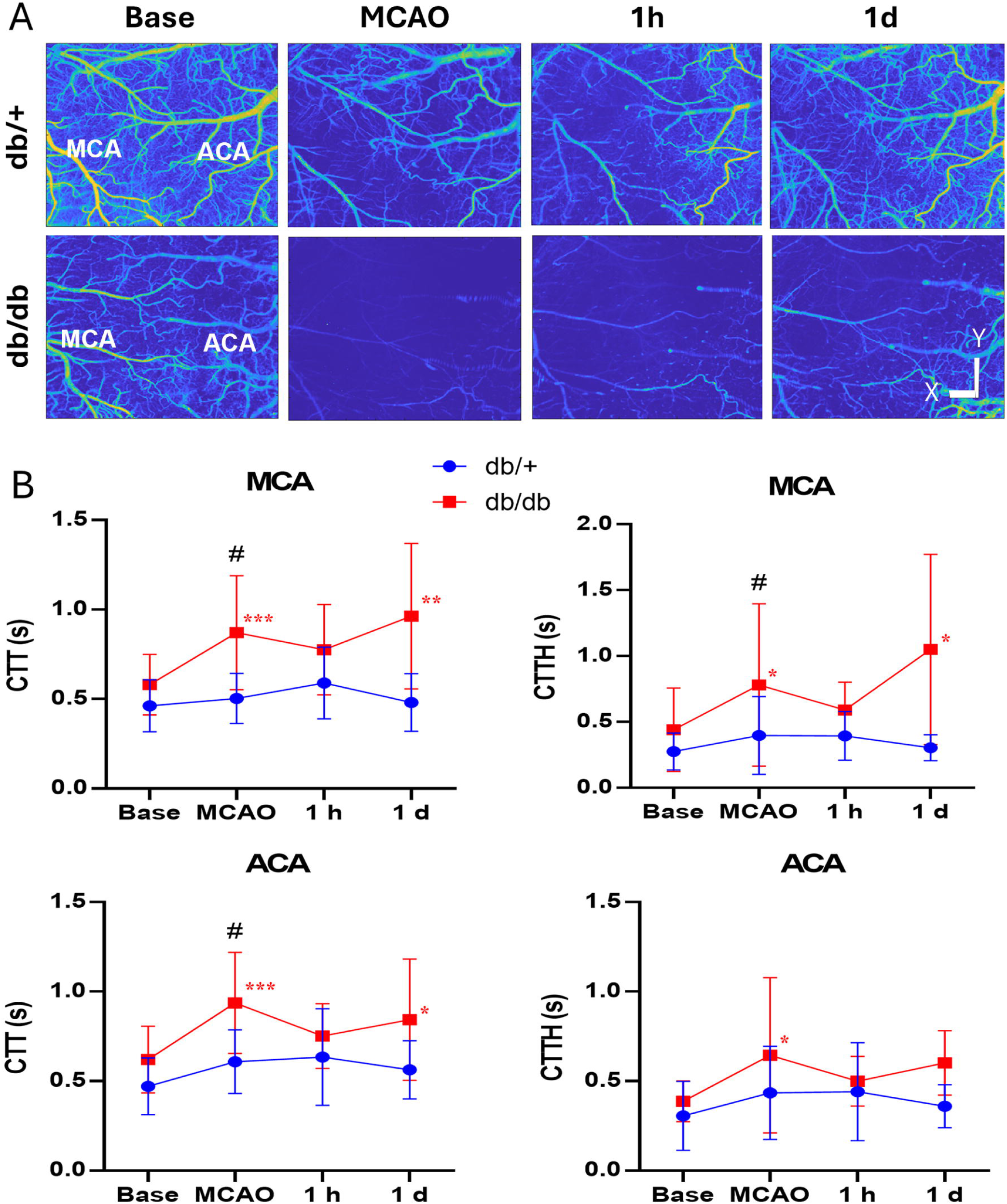
Diabetic mice displayed a greater spatial heterogeneity in capillary transit time (CTT) following stroke compared to db/+ mice. (A) Representative capillary mean frequency maps generated by *en face* projection of 3D frequency signal within cortical slabs for each time point as indicated. Scale bar represents 1 mm for X/Y dimensions. (B) A significant increase in mean CTT occurred after MCAO and extended till 1d in the db/db mice in both MCA and ACA territories. Capillary transient time heterogeneity (CTTH) increased after stroke and decreased after reperfusion but elevated again in the MCA territories in db/db mice. Time points vs. baseline: *, **, *** P < 0.05, 0.01, 0.005 (db/db). Strain difference: #, ##, ### P < 0.05, 0.01, 0.005. N = 14 db/+, 17 db/db.

## 4. Discussion

T2DM is associated with poor cerebral perfusion after acute ischemic stroke, yet the mechanism underlying the poor blood flow and the interaction between immune cells and endothelium in the neurovascular unit are not well understood. Using two-photon imaging, we demonstrated that the diabetic mice exhibited sustained poor leptomeningeal retrograde collateral flow following permanent MCAO compared to their normoglycemic counterpart. The deficit in flow compensation in the pial collaterals also propagated to a persistent reduction of blood flow in the penetrating arterioles in the distal MCA and ACA territories in these mice as demonstrated by OCT imaging. Furthermore, the recovery of retrograde flow velocity and flux of MCA branches were only seen in db/+ mice over seven days after MCAO, leading to a favorable cortical perfusion as reflected by the microvascular velocimetry data in CTT and CTTH. Most noticeably, distal arteries and veins of the diabetic mice are smaller in sizes prior to stroke, suggesting that T2DM chronically remodeled the leptomeningeal circulation, underlying the poor collateral flow seen after stroke. These results collectively support the poor collateral circulation between MCA and ACA and possibly an impaired clearance of inflammatory cell aggregates in the T2DM db/db mice. Our data also showed that the diabetic mice had persistently high level of leukocyte-endothelium adhesion in the cerebral veins, coincided with an increased number of neutrophils infiltrated into brain parenchyma after stroke compared to control mice, which may potentially contribute to a more severe ischemic injury in these mice.

To the best of our knowledge, this is the first study examining hemodynamics and leukocyte adhesion following ischemic stroke in awake condition using two-photon imaging. Anesthetics are commonly used in experimental stroke blood flow imaging studies due to technical necessity. However, anesthesia has known effect on blood vessel and neurons, thus can confound the results of hemodynamics and neuroprotection. A previous study using the photothrombotic stroke method comparing the neuroprotective effect of GluN2B and a4b2 nicotinic receptor antagonists found if stroke induced in awake mice, there was no neuroprotection by either drug in contrast to their reported benefit in reducing ischemic damage ^31^. Prior studies of two-photon were conducted under isoflurane, chloral hydrate medetomidine, midazolam or fentanyl, which may affect cerebral autoregulation, neurovascular coupling and vasomotor reactivity^32^. In this study the effect of anesthesia is exempt, allowing us to determine the effect of ischemic stroke on blood flow dynamics and the movement of blood cells under physiological stroke without the influence of anesthesia.

Experimental and clinical data suggest that T2DM is associated with poor collateral status during acute ischemic stroke^11, 12, 33^. However, the mechanism that led to the impaired retrograde compensation of pial vessel after ischemic stroke is not entirely clear. In the previous study, we found that the db/db young adult mice had a similar number of connecting collaterals at baseline, but a lower number of perfused collaterals at day seven after MCAO compared with db/+ mice ^11^. This result indicates that there is a functional impairment of collateral recruitment rather than the structural deficiency in the collateral circulation in db/db mice. One potential mechanism contributing to the functional impairment of collateral recruitment is an impaired endothelial regulation of vascular tone through nitric oxide (NO), which dilates the vessels and inhibits platelet activation^34^. Inhalation of NO during ischemia promotes cerebral collateral recruitment and reduces infarct size^35^. Conversely, reduced production of NO via impaired endothelial NO synthase (eNOS) decreased collateral flow recruitment and enlarge infarct sizes ^36^. Hyperglycemia inhibits the eNOS and thus cause the reduction of NO^37^, while restoring eNOS activity improves CBF in db/db mice^38^. Furthermore, plasma concentration and activity of plasminogen activator inhibitor-1, a potent endogenous tPA inhibitor, are heightened in T2DM^39, 40^. This might lead to increased formation of occlusive thrombi after vascular injury and thus could in part contribute to the impaired collateral status seen in the diabetic mice during ischemic stroke. Via two photon imaging, we found that the distal MCA branches maintaining anterograde flow after MCAO showed a dramatic decrease in velocity and flux in both genotypes compared to the MCA branches with retrograde flow. Furthermore, the velocity and flux of S1 and S2 with the anterograde flow were significantly lower compared to the MCA branches with retrograde flow in both genotypes. It suggests that the MCA branches without direct collateral from the ACA branches had a less favorable perfusion and a low ability to recover after stroke. These results suggest the importance of not only the functional collateral recruitment but the needed structural connection between ACA and MCA for perfusion and recover after stroke. Not surprisingly, therapies by enhancing collateral circulation are known to improve stroke outcome ^41–43^.

In the present study venous flow velocity in db/db mice also reduced persistently after MCAO compared to db/+ mice. In addition, db/db mice showed a progressive increase of WBC adhesion in the veins immediately after MCAO to at least 7ds afterwards. T2DM is associated with high basal cytokine levels and a wide range of innate immune responses ^44^, through which hyperglycemia has been linked to a pro-inflammatory state leading to increased production of interleukin-6 and tumor necrosis factor α ^45, 46^. Under the condition of ischemic stroke, peripheral myeloid cells including neutrophils are captured near the inflammatory site via rolling or adhesion on the brain vessel wall before transmigrating out of the bloodstream, as mediated by intracellular adhesion molecule 1 (ICAM-1), macrophage-1 antigen (MAC-1), and selectins ^47, 48^. In a mouse MCAO stroke model, genetic deletion of ICAM-1 reduces infarct volume, decreases mortality, and improves outcomes ^49^, although antibody targeted against ICAM-1 (Enlimomab) failed to provide neuroprotection in clinical trial ^50^. Neutrophil adhesion is also mediated by the integrin MAC-1. knocking out or inhibition of MAC-1 reduces the infarct size, mortality and neutrophil infiltration into the ischemic brain ^51^. Similarly, the success of MAC-1 inhibition in animal ischemic stroke also led to two ill-fated human studies ^52, 53^. Numerous studies have investigated the relationship between neutrophil adhesion and P-selectin during ischemic stroke. One study showed that soluble P-selectin levels were significantly higher in ischemic stroke patients than in control patients and the levels continued to rise until the subacute phase ^54^. In another study, the elevation of sodium P-selectin elements were linked to the progression of ischemic stroke ^55^. However, therapies targeting P-selectins in human ischemic patients have not shown benefit. Using a novel fusion peptide with P-selectin blocking and complement inhibitory activity (2.12Psel-Crry), we found that it inhibited leukocyte rolling after stroke, although the effect was not long lasting in the db/db mice ^56^.

The exact relationship between neutrophil and poor collateral circulation in T2DM after ischemic stroke is unclear; however, immune responses to ischemic stroke and adhesion of immune cells to endothelium may have profound effects on the flow dynamics. Studies from fluid dynamics of the blood suggest that shear- and conduit-dependent viscosity of blood cells may alter flow resistance in the vasculature via interaction among blood cells themselves, between blood cells and vessel wall, as well as between blood cells and fluid ^57, 58^. Due to the greater abundance of numbers in the blood, RBCs are known to exclude WBCs from the bulk flow, inducing leukocyte margination ^59^. Clumping of RBCs due to microthrombi formation following ischemic stroke especially increases the force of pushing WBC to the vessel wall, thus increasing the propensity of rolling and adhesion of the latter. T2DM is suggested to exacerbate p-selectin endothelial expression and also increase hematocrit and blood viscosity, thus further enhances rolling and adhesion of leukocytes after stroke ^60, 61^. The unique combination of vessel geometry and shear-dependent flow anomalies in post venule capillaries tends to promote RBC re-organization and aggregating, further increasing WBC rolling and adhesion ^62^. WBCs, albeit lesser in number yet with a larger size, can also affect the flow behavior of RBCs and flow dynamics. Finally, platelet activation induced by stroke and T2DM, further compounds the formation of cell aggregates and contributes to increased flow resistance and reduced flow rate ^63^. Flow stagnation is frequently observed in capillaries after stroke due to dynamic leukocyte stalls. By using a transgenic mouse model with tdTomato-expressing neutrophils, one study elegantly identified repetitively occurring capillary stalls of varying duration promoted by neutrophils in the permanent stroke models ^64^. It indicates that neutrophils play a major role in leukocyte rolling, adhesion, capillary stalling and flow obstruction. Prior studies have shown that patients with T2DM exhibit immune dysfunction related to neutrophil adhesion ^65, 66^, and that neutrophils taken from T2DM patients had a decreased rolling velocity than a healthy control population ^67^, suggesting that T2DM might increase neutrophil extravasation following ischemic stroke. In support of these reports, our FACS data showed that there were more neutrophils infiltrated into brain parenchyma in the db/db mice after stroke compared to db/+ mice.

In addition, neutrophils promoted-thrombosis may further exacerbate vessel occlusion and reduction of CBF following stroke. Emerging evidence indicates that neutrophil is a key contributor for thrombosis ^68^. Thrombus are promoted by neutrophils through several participants and mechanisms including tissue factor, interactions with platelets, release of Neutrophil Extracellular Traps (NETs), and release of proteases ^69, 70^. Activated neutrophils are an important source of tissue factor which interacts with coagulation factor VIIa, activates the extrinsic coagulation pathway, leading to thrombus formation ^70^. Neutrophils can also interact with platelets that result in enhanced platelet aggregation and clot formation ^71^. Neutrophil–platelet complexes are increased in patients with ischemic stroke and several drugs are indicated to prevent thrombus ^72, 73^. Derived from neutrophils and composed primarily of DNA, NETs are suggested to trigger platelet activation and promote thrombus formation ^74^. In addition, neutrophil proteases and thrombosis neutrophil-derived proteases also contribute to thrombus formation. Cathepsin G and neutrophil elastase act on coagulation factor X to promote coagulation ^75^. As such, antiplatelets and anticoagulation therapy is the gold standard treatment in reducing thrombosis in acute ischemic stroke ^76^. Clopidogrel (CPG) is one of the major antiplatelets used for ischemic stroke medication by inducing confirmation change in gpIIbIIIa, preventing neutrophil–platelet interaction and blocking platelet activation ^73, 76^. In a recent trial evaluating the efficacy of CPG on post stroke flow dynamics in the db/db and db/+ mice, we unexpectedly found no benefit of CPG treatment on CBF improvement nor rolling reduction in either type of mice (unpublished observation). One potential cause for the failure of CPG in improving blood flow is attributed to the permanent MCAO mode used in our study. In reported studies targeting neutrophil during ischemic stroke, beneficial effect was mostly observed in transient MCAO models, although the mechanism underling reperfusion-mediated neuroprotection by CPG is unclear ^49, 77^. Further studies are needed to evaluate the benefit of CPG on improving hemodynamics and reducing neutrophil-promoted thrombus formation during ischemic stroke comparing the effect of reperfusion.

One main limitation of our study lies in the animal model of T2DM. Leptin signaling plays a crucial role in developing obesity and insulin resistance, the genetic model db/db mice with *lepr* mutation was developed to model the leptin signaling deficiency in T2DM. Apart from being hyperglycemic, many metabolic and pathophysiological features exhibited in db/db mice are consistent with T2DM patients such as reduced adiponectin level, chronic inflammation, brain atrophy, bone fragility and poor stroke outcome including poor cerebral collateral flow or overall cerebral perfusion ^11, 12^. Recently we found that db/db mice also exhibited abnormal features of brain oscillations ^78^, similar to some reported in T2DM patients. However, db/db model does have limitations in modeling a complex human metabolic disease. First, although single-nucleotide polymorphism of LEPR was reported to be associated with T2DM ^79^, per CDC most T2DM humans fell sick mainly due to their diets, instead of their genetics (CDC diabetes statistics). Another pitfall of the db/db mice is that T2DM patients may have increased, decreased or neutral level of leptin compared to non-diabetic humans, unlike the high leptin level in the db/db mice. Besides, the juvenile onset of hyperglycemia in the db/db model is not consistent with the human T2DM disease process. Disrupting leptin signaling at an early stage may affect many aspects of development. Thus, a diet model such as C57BL/6 mice fed with high fat diet (HFD) is more sensible and became the model of choice for diet-induced obesity pre-clinical stroke studies including SPAN ^80, 81^. Interestingly, despite the rational consideration of the HFD model, one study pointed out that db/db mice captured some key features of human T2D pancreatic islet cell pathology that are not present in weight-matched C57BL/6J mice fed a western diet, such as increased glucagon, increased islet expression of the dedifferentiation marker Aldh1a3, or reduced nuclear presence of the transcription factor Nkx6.1, suggesting that in body-weight matched mice, the WD-fed C57BL/6J mouse is an excellent model of early features representing the human prediabetic condition, while the db/db mouse more closely resembles human T2D ^82^. The unexpected smaller infarct found in mice fed with HFD compared to those with normal diet from the SPAN trial was also not in line with human stroke patients with T2DM ^83^. The inconsistent pathophysiology between human and animals with T2DM further bolsters the limitations of animal models of diabetes ^84–86^. Nevertheless, it would be important to be able to confirm the dysregulation of cerebral flow and immune cell adhesion to endothelium in the HFD model as seen in the db/db model. A second major limitation of the current study is the use of permanent MCAO model for evaluating the dynamics of retrograde pial flow after stroke due to the growing success of recanalization therapy in large artery strokes. It is likely that retrograde leptomeningeal blood flow from ACA will dissipate and anterograde flow from MCA restored. Nevertheless, there is still a significant fraction of brain vessels remain unperfused either due to other stroke subtypes or incomplete recanalization. Lastly, due to the limitation of our study approach, we cannot characterize how each cell type in the neurovascular unit affected by T2DM after stroke. It is well known that astrocytic foot processes were detached and microglial cells were indicated cause invasive damaging during the injury in diabetic mice ^87^. Pericyte apoptosis has also been recognized in vivo in hyperglycemic rats and in vitro in retinal cultures ^88^. Furthermore, endothelial cell deterioration at the electronic-microscopic level was observed in diabetic mice including the loss of electron density, basement membrane thickening and rearrangement, increased transcytotic-pinocytotic vesicles, and aberrant mitochondria ^1^. The data obtained from the current study opens up a rich venue to determine the underlying pathological mechanism leading to the poor perfusion and poor stroke outcome in human T2DM.

## 5. Conclusion

Compared to control mice, T2DM mice displayed persistently reduced blood flow in pial arteries and veins, coincided with a poor recovery of penetrating arterial flow and sustained deficit in microvascular flow. There was also persistent increase of leukocyte adhesion to endothelium of veins and elevated neutrophils infiltration into brain parenchyma in T2DM mice compared to control mice after stroke. Taken together, our data suggest that T2DM induced increase in inflammation and chronic remodeling of leptomeningeal vessels may contribute to the observed hemodynamics deficiency and poor stroke outcome.

## Author Contribution Statement

Yoshimichi Sato: Writing – original draft, Visualization, Methodology, Investigation, Formal analysis, Data curation, Conceptualization. Yuandong Li: Methodology, Software and analysis of PA flow. Yuya Kato: Investigation, Data curation. Atsushi Kanoke: Investigation, Data curation of OCTA. Jennifer Y Sun: Methodology, Software of two-photon imaging. Yasuo Nishijima: Investigation, Data curation of FACS. Ruikang K. Wang: Methodology, Software of OCTA. Michael Stryker: Methodology, feasibility, Software of two-photon imaging. Hidenori Endo: insights of clinical relevance. Jialing Liu: Writing – review & editing, Visualization, Supervision, Project administration, Methodology, Investigation, Funding acquisition, Formal analysis, Data curation, Conceptualization.

## Ethics Statement

The study was conducted in accordance with the Guide for Care and Use of Laboratory Animals issued by the National Institutes of Health and approved by San Francisco Veterans Affairs Medical Center Institutional Animal Care and Use Committee (protocol number 20-012, approved 8-6- 2020).

## Acknowledgements

This work was supported by NIH grant R01NS102886 (JL), VA Merit Award 2I01BX003335 (JL), and Research Career Scientist award 2IK6BX004600 (JL).

## Abbreviations

T2DM: Type 2 diabetes mellitus
OCT: Optical Coherence Tomography
MCAO: Middle Cerebral Artery Occlusion
MCA: Middle Cerebral Artery
ACA: Anterior Cerebral Artery
PA: Penetrating Artery
MF: Mean Frequency
CTT: Capillary Transit Time
CTTH: Capillary Transit Time Heterogeneity
NO: Nitric Oxide
eNOS: endothelial Nitric Oxide Synthase
ICAM-1: intracellular adhesion molecule 1
MAC-1: intracellular adhesion molecule 1

