## Supplemental Materials for "Type 2 diabetes remodels collateral circulation and promotes leukocyte adhesion following ischemic stroke"


This PDF file includes:

- Supple Method for OCT
- Supple Figure S1

Supple Tables S1 to S2

Measurement of cerebral blood flow dynamics via optical coherence tomography angiography (OCTA)

System configuration

Cortical penetrating arterial microvascular flow dynamics were quantified by a custom-built spectral domain OCT system equipped with a broadband super-luminescent diode (SLD) light source (LS2000B, Thorlabs Inc.) with a center wavelength of 1310 nm and a spectral bandwidth of ~110 nm at 3dB, providing an axial resolution of ~6 µm in tissue as previously described [11, 12, 21] (Supplemental Figure 1A). High-resolution OCT imaging was conducted through a cranial window using a 10X lens and a visible light camera was coupled to the system to facilitate targeting the scan area. The light beam was scanned over the cranial window using a paired X-Y galvanometer (6210, Cambridge Technology), yielding a 3-D volumetric dataset (z-x-y). The backscattered signal interfering with the counterpart from the reference arm is detected by a spectrometer consisting of a transmission grating, an achromatic doublet lens, and a 1024-pixel InGaAs line scan camera operated at 92,000 axial scans per second. The system sensitivity was measured to be 105 dB at the focus (~500 µm below the zero-delay line). A total of four temporal scans were performed under isoflurane anesthesia for each imaging modality below: baseline prior to stroke (labeled as Base), 30 minutes after the occlusion of MCA and ipsilateral CCA (as MCAO), 1 hour after the reversal of CCAO (as 1 h), and 1 day after MCAO (as 1 d).

OCT microangiography (OMAG)

Morphological features of functional cerebral blood flow were obtained via the OMAG scanning protocol, consisting of 400 A-lines per B-scan in the x axis, multiplied by 400 clusters of 8 repeated B-scans per cluster in the y-axis. This protocol was performed over 9 tiles facilitated by a motorized translational stage to cover the desired field of view. The total scan time for OMAG was 5 min, generating a final 3-D OMAG data of 400 x 400 x 512 (x-y-z) voxels. The flow signals extracted using our custom OMAG algorithm accompanied by blood flow maps were shown as the en face (x-y) maximum intensity projection of the 3-D volume (Supplemental Figure 1B).

Doppler optical microangiography (DOMAG)

DOMAG was used to quantitatively evaluate the flow dynamics of the penetrating arterioles (PA) in the distal MCA and ACA territories. The DOMAG scanning protocol consists of 25 A-lines repeatedly acquired to form an M-scan (z axis), multiplied by 300 B scans on the y-axis at 380 M-scan steps per B scan on the x-axis, producing a 3-D data set of 380x300x512 (x-y-z) voxels. The total scan time of DOMAG was 20 min. The directional blood flow velocity (axial component) was deduced by the positive or negative phase changes calculated between A-line intervals (Supplemental Figure 1C), and the velocity signals were false color-coded to represent two opposite flow directions with penetrating arterioles in green and the rising venules in red in the final en face image (Supplemental Figure 1D). A wide range of flow velocity can be obtained with this algorithm through A-line skipping, and the displayed bidirectional velocity maps in this study had a flow range of ± 6.1mm/s. The DOMAG scanning was performed over 4 tiles to produce the final flow velocity map of 3 mm x 2 mm/tile. The identification of true PA signals was aided by 3-D data visualization in Amira (Zuse Institute Berlin and Thermo Fisher Scientific) and confirmed with structural images, and an x-y orthoslice at 75 µm below the surface (Supplemental Figure 1E) showing the cross section of the flow velocity signals was selected for quantification. The encircled signals carry information of flow cross-sectional area ($mm^2$), flow velocity (mm/s), and flow rate ($mm^3$/s) within MCA and ACA regions

(Supplemental Figure 1F), which can be calculated based on the method previously described [21, 26].

OCTA capillary velocimetry

The capillary flow dynamics was assessed using a OCTA capillary velocimetry method described previously [27], in which A-lines were repeated 50 times to form an M-scan complex with a total scan time of 20 min. The A-line speed was set at 20 kHz with 50 repeats to yield an A-line interval between 50 µs to 2.5 ms, which could potentially capture fast to slow capillary velocity from 5 mm/s to 100 µm/s. A B-frame consisted of 400 M-scans, and there were 400 B-frames in the 3-D dataset. The final 3-D cube had 512 x 400 x 400 voxels covering a 4 mm x 4 mm region. A special mask was additionally applied to B-frames to remove vessels with lumen larger than 15 µm to achieve better visualization and quantification of the mean frequency (MF) signals from capillaries (Figure 2G). A set of en face average intensity projections (AIP) within a 300 µm-thick slab was made to visualize the capillary MF maps within the scanning region (Supplemental Figure 1H). A statistical method of Eigendecomposition (ED) was performed where the eigenvalues representing static signals were removed and the mean frequency (MF) of moving particles was estimated. Then, the MF signals were converted to velocity according to the linear relation derived from the previous phantom experiment in microfluidics[27]. To obtain transit time parameters, the velocity is converted to time by dividing mean capillary path length of 400 mm as previously described [28]. Capillary transit time distribution is expressed in the gamma function as $h(\tau;\alpha,\beta) = \frac{1}{\beta^{\alpha}\Gamma(\alpha)}\tau^{\alpha-1}e^{-\tau/\beta}$ [28, 29]. Finally, the mean capillary transit time (CTT) and capillary transit time heterogeneity (CTTH) were calculated by the mean (ab) and standard deviation ($\sqrt{\alpha}$b) from the gamma function (Supplemental Figure 1I).

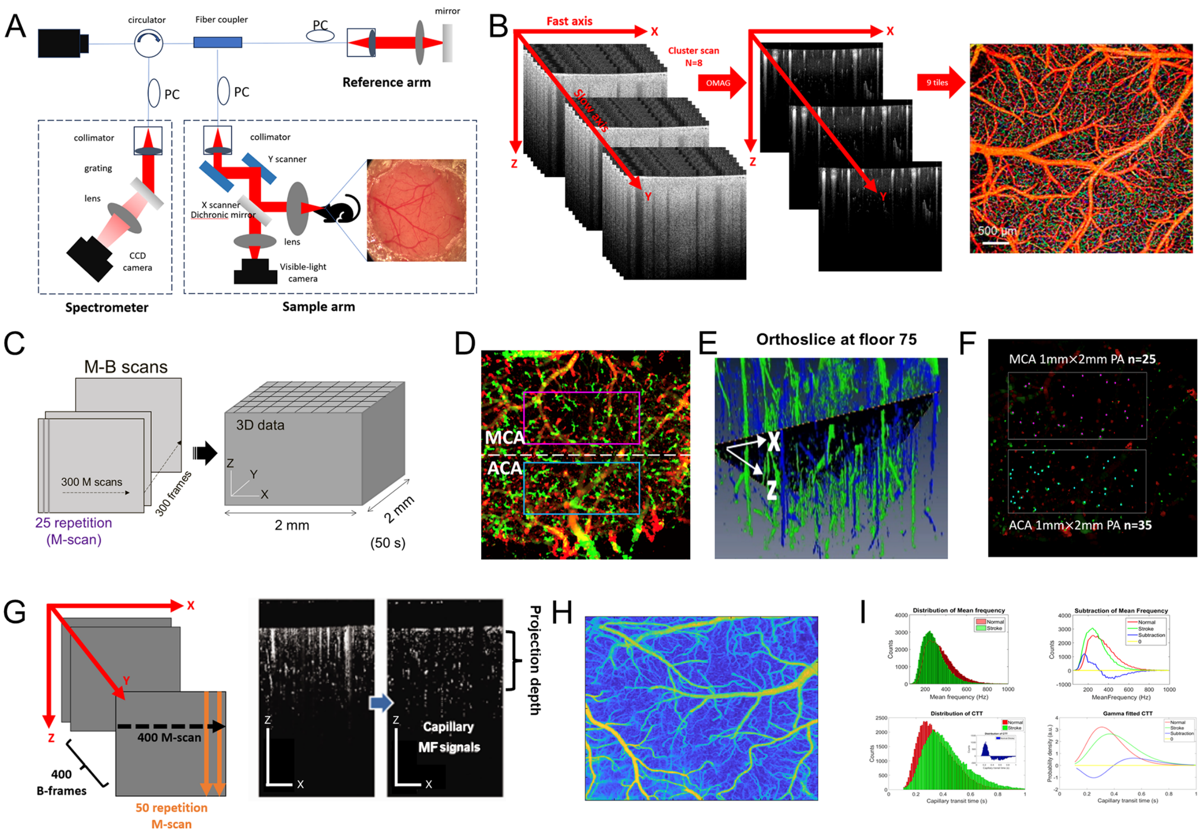

**Supplemental Figure 1. The schematic diagram of the optical coherence tomography (OCT) system and methods to determine penetrating arterial and microvascular flow.** (A) The schematic of the custom-designed spectral domain OCT system using a 1310 nm light source. (B) Optical microangiography (OMAG) scanning scheme. Eight repeated scans were performed in one cluster, where OMAG algorithm was applied to extract blood flow information. A total of 400 clusters of scans (400 frames) was obtained to form one OMAG volume. Nine OMAG volume image were then stitched to generate one *en face* OMAG maximal projection image. (C) The scheme of the DOMAG method scanning protocol. 25 A-lines repeatedly acquired to form an M-scan (z axis), multiplied by 300 B scans on the y-axis at 380 M-scan steps per B scan on the x-axis, producing a 3-D data set of 380x300x512 (x-y-z) voxels. (D) Velocity map of a maximal intensity projection of the penetrating arteries (green) and penetrating veins (red) and areas of quantification. (E) demonstration of a X-Y plane at floor 75 of the orthoslices from the 3D dataset. (F) 25 and 35 PAs were identified in the 1mm x 2mm grids of MCA and ACA territories, respectively, and flow velocity was calculated. (G) Schematics of OCTA velocimetry scanning protocol. Mask was applied to MB scans to remove signals from vessels with lumens larger than 15 µm. (H) Capillary mean frequency maps generated by *en face* projection of 3D frequency signal within a 300 um-thick slab. (I) Distribution of mean frequency and capillary transit time at baseline (Normal) and 30 minutes after dMCAO (Stroke), showing the difference between two conditions. Difference in mean frequency between two conditions was shown as the subtracted value, and CTT was calculated using the gamma function.

| Main Effects | genotype | timepoint | interaction |
|---|---|---|---|
| **Velocity** | | | |
| S1 revers | **< 0.05** | **< 0.0001** | **< 0.0001** |
| S2 revers | 0.1232 | **< 0.0001** | **< 0.0001** |
| S3 revers | 0.3954 | **< 0.0001** | **< 0.01** |
| vein | **< 0.0001** | **< 0.0001** | **< 0.0001** |
| S1 anterograde | 0.2588 | **< 0.0001** | 0.1132 |
| S2 anterograde | 0.5799 | **< 0.0001** | 0.235 |
| **Diameter** | | | |
| S1 revers | 0.3255 | **< 0.0001** | **< 0.0001** |
| S2 revers | 0.9792 | **< 0.0001** | 0.4852 |
| S3 revers | 0.0533 | **< 0.0001** | 0.68785 |
| vein | **< 0.01** | **< 0.0001** | **< 0.05** |
| S1 anterograde | 0.5282 | **< 0.0001** | **< 0.05** |
| S2 anterograde | 0.1256 | **< 0.0001** | 0.6387 |
| **Flux** | | | |
| S1 revers | **< 0.0001** | **< 0.001** | 0.7699 |
| S2 revers | **< 0.0001** | **< 0.0001** | 0.1166 |
| S3 revers | 0.0587 | **< 0.0001** | 0.647 |
| vein | **< 0.0001** | **< 0.0001** | **< 0.01** |
| S1 anterograde | 0.321 | **< 0.0001** | **< 0.05** |
| S2 anterograde | **< 0.05** | **< 0.0005** | 0.9854 |
| **Vein** | genotype | segment | interaction |
| Velocity | **<0.001** | **<0.01** | **<0.05** |
| Diameter | **<0.001** | **<0.001** | 0.4145 |
| Flux | **<0.001** | **<0.001** | **<0.005** |
| Leukocyte adhesion | **<0.01** | 0.1324 | 0.1523 |

**Supplemental Table 1**. Summary of p values of the two-way ANOVA test. Main effect of genotype, timepoint and interactions among velocity, diameter and flux for S1, S2, S3, veins, S1 and S2 with anterograde flow are described. Values less than 0.05 are demarked in bold. Significant main effect of genotype (velocity: S1 $p < 0.05$, Vein $p < 0.0001$, S1 anterograde < 0.0005; diameter: Vein $p< 0.01$ ; flux: S1 $p < 0.0001$, S2 $p < 0.0001$, Vein $p < 0.0001$, S1 anterograde < 0.01, S2 anterograde < 0.01) and timepoint (velocity: S1 $p < 0.0001$, S2 $p < 0.0001$, S3 $p < 0.0001$, Vein $p < 0.0001$, S1 anterograde $p < 0.0001$, S2 anterograde $p < 0.0001$ ; diameter: S1 $p < 0.0001$, S2 $p < 0.0001$, S3 $p < 0.0001$, Vein $p < 0.0001$, S1 anterograde $p < 0.0001$, S2 anterograde $p < 0.0001$; flux: S1 $p < 0.001$, S2 $p < 0.0001$, S3 $p < 0.0001$, Vein $p < 0.0001$, S1 anterograde $p < 0.0001$, S2 anterograde $p < 0.0001$) was recognized. Additionally, a significant interaction between genotype and timepoint was observed in some MCA segments and veins for velocity (S1 $p < 0.0001$ S2 $p < 0.0001$, S3 $p < 0.01$, Vein $p < 0.0001$, S2 anterograde $p < 0.0001$), diameter (S1 $p < 0.0001$, Vein $p < 0.05$, S1 anterograde $p < 0.01$) and flux (Vein $p < 0.01$), indicating that the effect of genotype varied across timepoints.

| | genotype | segment | interaction |
|---|---|---|---|
| Velocity | **<0.001** | **<0.01** | **<0.05** |
| Diameter | **<0.001** | **<0.001** | 0.4145 |
| Flux | **<0.001** | **<0.001** | **<0.005** |
| WBC adhesion | **<0.01** | 0.1324 | 0.1523 |

**Supplemental Table 2**. Summary of two-way ANOVA test. Main effect of genotype, vein segment and interaction among velocity, diameter and flux. Values less than 0.05 are demarked in bold. Significant differences were found in the main effect of genotype in velocity ($p<0.0001$), diameter ($p<0.0001$), flux ($p<0.0001$) and leukocyte adhesion ($p<0.01$). Significant differences were also found between the vein segments in velocity ($p<0.01$), diameter ($p<0.0001$), and flux ($p<0.0001$), but not in leukocyte adhesion.
